# Cyclin B3 promotes APC/C activation and anaphase I onset in oocyte meiosis

**DOI:** 10.1101/390161

**Authors:** Mehmet E. Karasu, Nora Bouftas, Scott Keeney, Katja Wassmann

**Author notes:** These authors contributed equally (Mehmet E. Karasu, Nora Bouftas). Corresponding authors: Katja Wassmann, Scott Keeney.

## Abstract

As obligate kinase partners, cyclins control the switch-like cell cycle transitions that orchestrate orderly duplication and segregation of genomes. Meiosis, the cell division that generates gametes for sexual reproduction, poses unique challenges because two rounds of chromosome segregation must be executed without intervening DNA replication. Mammalian cells express numerous, temporally regulated cyclins, but how these proteins collaborate to control meiosis remains poorly understood. Here, we delineate an essential function for mouse cyclin B3 in the first meiotic division of oocytes. Females genetically ablated for cyclin B3 are viable, indicating the protein is dispensable for mitotic divisions, but are sterile. Mutant oocytes appear normal until metaphase I but then display a highly penetrant failure to transition to anaphase I. They arrest with hallmarks of defective APC/C activation, including no separase activity and high MPF, cyclin B1, and securin levels. Partial APC/C activation occurs, however, as exogenously expressed APC/C substrates can be degraded and arrest can be suppressed by inhibiting MPF kinase. Cyclin B3 is itself targeted for degradation by the APC/C. Cyclin B3 forms active kinase complexes with CDK1, and meiotic progression requires cyclin B3-associated kinase activity. Collectively, our findings indicate that cyclin B3 is essential for oocyte meiosis because it fine-tunes APC/C activity as a kinase-activating CDK partner. Cyclin B3 homologs from frog, zebrafish, and fruitfly rescue meiotic progression in cyclin B3-deficient mouse oocytes, indicating conservation of the biochemical properties and possibly cellular functions of this germline-critical cyclin.

## Introduction

Eukaryotic cell division depends on oscillations of cyclin-dependent kinases (CDKs) associated with specific cyclins [1–4]. In vertebrate somatic cells, progression from G1 into S phase, G2, and mitosis depends on the cyclin D family, followed by the cyclin E, A, and B families [4]. Ordered CDK activity likewise governs progression through meiosis: chromosome condensation, congression, and alignment require a rise in cyclin B-CDK1 activity, then anaphase onset and chromosome segregation are driven by sudden inactivation of cyclin B-CDK1 by the anaphase promoting complex/ cyclosome (APC/C), an E3 ubiquitin ligase that targets substrates for degradation by the 26S proteasome [5]. Cyclin B-CDK1 activity re-accumulates following its depletion, but meiotic cells have the challenge of not inducing DNA replication during the period of low cyclin B-CDK1 activity between meiosis I and II [6, 7]. The roles of specific cyclins in shaping CDK oscillations and thus in determining the orderly progression of meiosis remain poorly understood, particularly in mammalian oocytes.

In this context, cyclin B3 is particularly enigmatic. Cyclin B3 belongs to a separate family with characteristics of both A- and B-type cyclins [8, 9] and is evolutionarily conserved from early metazoans to human [10]. *Ciona intestinalis* cyclin B3 counteracts zygotic genome activation, thus maternal cyclin B3 mRNA diminishes at zygotic transcription onset [11]. In *Xenopus laevis*, no cyclin B3 protein was detected by western blot in oocytes or during the first embryonic divisions [12], but whether cyclin B3 has a role in meiotic or embryonic cell cycle progression in the frog has not been addressed. *Drosophila melanogaster* cyclin B3 is dispensable for mitotic divisions and for male fertility, but is essential for female fertility [13]. In flies, cyclin B3 promotes anaphase onset in early embryonic divisions [14], and loss of cyclin B3 perturbs exit from meiosis I [13]. Moreover, *Drosophila* cyclin B3 is degraded depending on the APC/C in mitosis, with delayed timing relative to cyclin A and B [14, 15]. Unlike in *Drosophila* [14], in *Caenorhabditis elegans* embryos cyclin B3 was proposed to promote anaphase onset in mitosis through a role in the spindle assembly checkpoint (SAC) [16, 17]. *C. elegans* and *Drosophila* cyclin B3 associate with CDK1, and *in vitro* kinase activity was detected [13, 17].

Interestingly, cyclin B3 protein has increased sharply in size in placental mammals due to the extension of a single exon [10]. Cyclin B3 may therefore occupy additional, unknown roles during cell division in mammals. In mice, cyclin B3 was proposed to be required for meiotic recombination in male meiosis, because its mRNA is present early in meiotic prophase I [18, 19]. Prolonged expression of cyclin B3 in testis perturbed spermatogenesis, and cyclin B3 interacts with CDK2, although no associated kinase activity was detected [19]. In female mice, cyclin B3 was speculated to be required for meiotic initiation because its mRNA is substantially upregulated as oogonia cease mitotic proliferation and enter meiotic prophase I [20]. RNAi-mediated knock-down of cyclin B3 to ~30% of wild-type levels in cultured mouse oocytes perturbed progression through meiosis I [21]. However, the molecular basis of the inhibition of meiotic progression was not defined and off-target effects of the RNAi could not be excluded, so the function of cyclin B3 in oocyte meiosis remained unclear.

To recapitulate, cyclin B3 has been implicated in female meiosis and early embryogenesis in different organisms, but its function remains poorly understood, particularly in mammals. To clarify whether cyclin B3 is indeed required for mammalian oocyte meiosis and to gain insights into its potential role(s), we generated mice with a targeted mutation in *Ccnb3*, the gene encoding cyclin B3. Mice are viable and males are unexpectedly fertile without cyclin B3, and contrary to earlier predictions [20] cyclin B3-deficient oocytes are proficient at entering meiosis. Strikingly, however, we find that cyclin B3 is required for female meiosis I, and most oocytes cannot progress beyond metaphase I in its absence. As a result, *Ccnb3*-deficient female mice are sterile. Our data indicate that cyclin B3 fine-tunes APC/C activity towards meiotic substrates during oocyte meiosis I, at least in part as a partner enabling CDK1 catalytic activity, and that the biochemical function of cyclin B3 is conserved in vertebrates.

## Results

### Female mice are sterile after genetic ablation of Ccnb3

To determine the function of mouse cyclin B3, we generated a targeted mutation by CRISPR/Cas9-mediated genome editing. The *Ccnb3* locus, which spans 62 kb on the X chromosome (coordinates chrX:6,556,778–6,618,745, mm10 genome assembly), gives rise to a 4.1-kb mRNA that comprises 14 exons and encodes a protein of 157.9 kDa. We used a guide RNA to target the 3ʹ end of the 2.7-kb long exon 7, generating a mutant allele with a 14-bp deletion causing a frameshift and premature stop codon upstream of the cyclin box, which is encoded in exons 9–13 (**Figure 1A**). Cyclin B3 protein was readily detected in extracts of wild-type adult testes by immunoprecipitation/western blotting, but extracts from *Ccnb3^−/Y^* mutant testes failed to yield signal for either full-length cyclin B3 or the predicted truncation product that would lack the cyclin box (1090 amino acids long, ~123 kDa) (**Figure 1B**). These results suggest that the truncated form of the protein is unstable if it is made at all, and indicate that the mutation is a null or severe loss-of-function allele.

**Figure 1.**
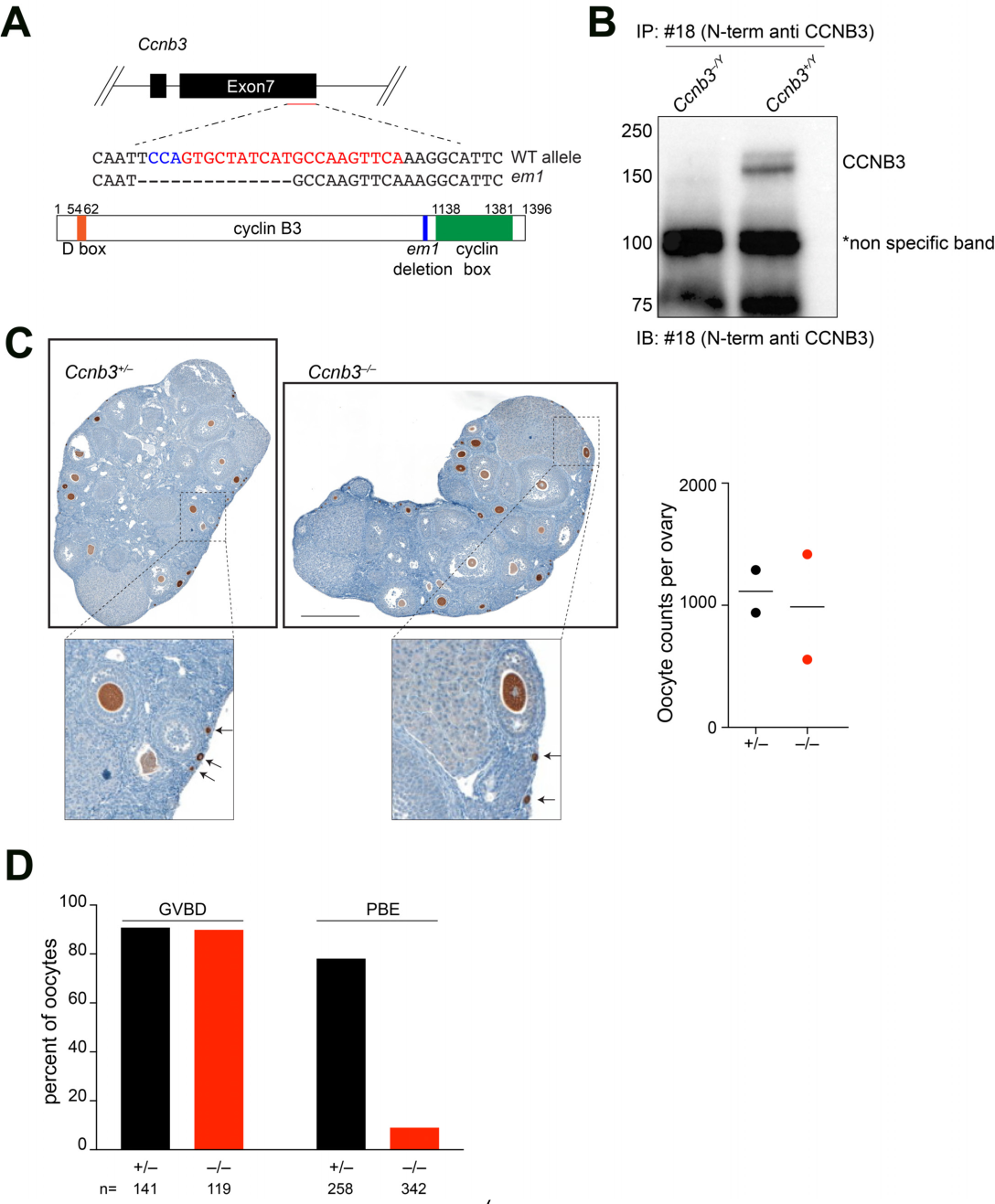
Generation of *Ccnb3^−/−^* mice reveals a requirement for cyclin B3 in female meiosis. (A) Targeted mutation of *Ccnb3*. (Top) Exons 6 and 7 are shown as black rectangles. A portion of the sequences of the wild-type allele and *em1* mutant allele are shown, with guide RNA position in red and the complement of the PAM sequence in blue. (Bottom) Schematic of cyclin B3 protein. (B) Immunoprecipitation and western blot analysis of adult testis extracts using anti-cyclin B3 monoclonal #18 antibody (Karasu et al., 2018, in preparation), which recognizes an N terminal epitope. (C) Apparently normal folliculogenesis and oocyte reserves in *Ccnb3*-deficient females. (Left) PFA-fixed, anti-MVH stained ovary sections from three-month-old animals. Zoomed images show presence of primary/primordial follicles indicated by black arrows. Scale bar represents 500 µm. (Right) Oocyte counts from two three-month-old females of each genotype. PFA-fixed ovaries were sectioned completely and stained with anti-MVH. Stained oocytes were counted in every fifth ovary section and summed. Each point is the count for one animal. (D) Meiosis I arrest. Percentages of mature oocytes (which were not maintained arrested in GV stage) that underwent GVBD (germinal vesicle breakdown) within 90 min in culture, and oocytes that extruded polar bodies (PB) (n: number of oocytes).

Surprisingly, despite prominent, regulated expression during spermatogenesis, cyclin B3-deficient males were fully fertile, with no detectable meiotic abnormalities (a detailed description of the male phenotype will be provided elsewhere). Heterozygous females also displayed normal fertility and homozygous mutant females were born in Mendelian frequencies from heterozygous parents (105 +/– (25.7%):100 –/– (25.5%):102 +/Y (25%):101 –/Y (24.8%) offspring from *Ccnb3^+/–^* females crossed with *Ccnb3^−/Y^* males). *Ccnb3^−/−^* animals displayed no gross abnormalities in histopathological analysis of major organs and tissues (see Methods), indicating that cyclin B3 is dispensable in most if not all non-meiotic cells.

Strikingly, however, cyclin B3-deficient females were sterile. No pregnancies were observed and no pups were born from *Ccnb3^−/−^* females (n=9) bred with wild-type males for two rounds of mating. We conclude that, whereas cyclin B3 is dispensable for male meiosis, it is essential for female fertility.

### Cyclin B3 is required for the metaphase-to-anaphase I transition in oocytes

To understand why *Ccnb3^−/−^* females are sterile, we examined the behavior of mutant oocytes. In mice, as in other mammals, female meiosis initiates during fetal development [6, 22]. Oocytes complete the chromosome pairing and recombination steps of meiotic prophase prior to birth, then enter a prolonged period of arrest (the dictyate stage) and coordinate with surrounding somatic cells during the first few days after birth to form follicles [6, 22]. Primordial follicles are the resting pool of germ cells that will be recruited for further development and ovulation during the reproductive life of the animal [6, 22]. Ovary sections from *Ccnb3^−/−^* females showed abundant oocytes in primordial and growing follicles at 2–3 months of age, and were quantitatively and morphologically indistinguishable from littermate controls (**Figure 1C)**. Follicle formation is highly sensitive to pre-meiotic and meiotic prophase I defects, such that mutations compromising oogonial development, meiotic initiation, or recombination result in drastically reduced or absent follicles within the first few weeks after birth if not earlier [23–27]. The normal number and appearance of follicles in *Ccnb3^−/−^* females therefore leads us to infer that cyclin B3 is largely if not completely dispensable for germ cell divisions prior to meiosis and for early events of meiotic prophase I (DNA replication, recombination, and chromosome pairing and synapsis).

We next tested the ability of mutant oocytes to be induced to enter meiosis I and to carry out the meiotic divisions *in vitro*. Germinal vesicle (GV) stage oocytes were collected from ovaries of adult *Ccnb3^−/−^* mice and age-matched controls from the same breedings (*Ccnb3^+/–^*) and cultured *in vitro*. Under these conditions, germinal vesicle breakdown (GVBD, corresponding to nuclear envelope breakdown in mitosis) occurred with normal timing (within 90 min) and efficiency in *Ccnb3^−/−^* oocytes **(Figure 1D)**, indicating that cyclin B3 is dispensable for entry into meiosis I. At ~7–10 hours in culture after entry into meiosis I, depending on the mouse strain, oocytes extrude a polar body (PB), indicating execution of the first division and exit from meiosis I [6, 28]. Unlike for other aspects of female reproduction described above, *Ccnb3^−/−^* mice displayed a highly penetrant defect at this stage of meiosis: oocytes failed to extrude a PB in most cyclin B3-deficient mice (50 out of 58 mice) (**Figure 1D**). These findings indicate that cyclin B3 is required for progression through meiosis I, suggesting in turn that meiotic progression failure is the cause of female infertility. Because the penetrance of meiotic arrest was mouse specific, (i.e., most mice yielded only arrested oocytes) we focused subsequent experiments on the large majority of mice where no PB extrusion was observed (see Methods). Possible reasons for the incomplete penetrance are addressed in the Discussion.

To further characterize this meiotic arrest, we followed progression through meiosis I by live imaging. GV oocytes were injected with mRNA encoding histone H2B fused to RFP and β-tubulin fused to GFP to visualize chromosomes and spindles, respectively. Whereas control oocytes separated their chromosomes and extruded PBs ~8 hours after GVBD, *Ccnb3^−/−^* oocytes that did not extrude PBs remained in a metaphase I-like state, with a spindle that had migrated close to the cortex and chromosomes aligned at the spindle midzone (**Figure 2A,** see **Movies 1 and 2** for chromosome movements).

**Figure 2.**
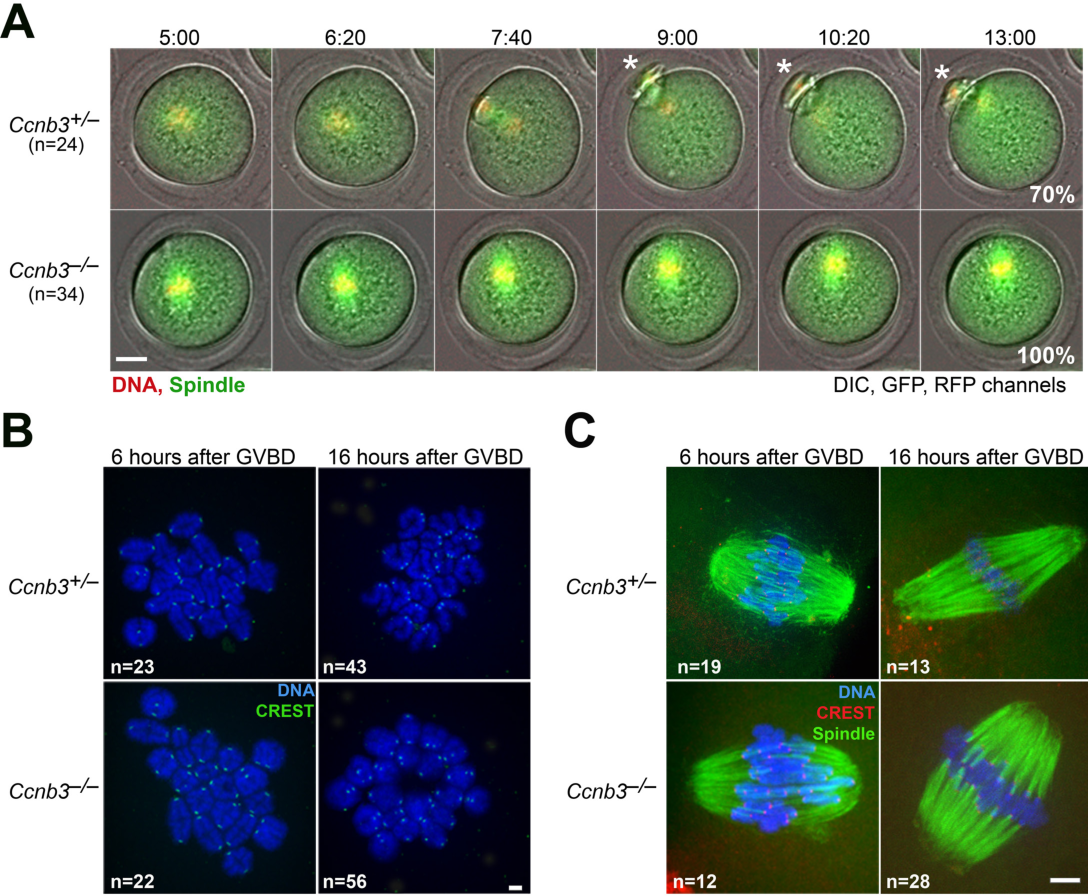
Cyclin B3 is required for the metaphase-to-anaphase I transition in oocytes. (A) Live-cell imaging of meiotic maturation. b-tubulin-GFP and H2B-RFP mRNA to visualize the spindle and chromosomes, respectively, were injected into GV-stage oocytes, which were then induced to enter meiosis I. Selected time frames are shown with an overlay of DIC, GFP and RFP channels from representative movies. Time after GVBD is indicated as hours:minutes. Percentage of oocytes of the observed phenotype is indicated. Scale bar represents 20 µm, white asterisks indicate PBs. Related to supplemental movie 1 and 2. (B) Chromosome spreads 6 hours (corresponding to metaphase I) and 16 hours (corresponding to metaphase II in controls) after GVBD. Kinetochores were stained with CREST (green) and chromosomes with Hoechst (blue). Scale bar represents 5 µm. (C) Whole-mount immunofluorescence staining of cold-treated spindles. Microtubules were stained with anti-tubulin antibody (green), kinetochores with CREST (red), and chromosomes with Hoechst (blue). Scale bar represents 5 µm.

We hypothesized that mutant oocytes were failing to make the transition from metaphase I to anaphase I; this idea predicts that homologous chromosomes will still be conjoined in those oocytes that failed to extrude PBs in the absence of cyclin B3. In chromosome spreads from control oocytes, we observed the expected 20 bivalent chromosomes (homolog pairs connected by chiasmata) prior to the first division (6 hours after GVBD) and pairs of sister chromatids juxtaposed near their centromeres afterwards (metaphase II arrest, 16 hours after GVBD) (**Figure 2B**). In contrast, as predicted, *Ccnb3^−/−^* oocytes displayed intact bivalents throughout the arrest in culture and no homolog separation was observed, indicating a metaphase I arrest (**Figure 2B**).

Arrest was not due to failure in forming either spindles or stable kinetochore-microtubule interactions, because metaphase I spindles displayed no obvious morphological defects and cold-stable microtubule fibers were observed in *Ccnb3^−/−^* oocytes comparable to control oocytes in metaphase I (**Figure 2C**). Microtubule fibers attached to kinetochores were clearly visible, indicating that cyclin B3 is not required for kinetochore-microtubule attachments (**Figure 2C**). Importantly, the bipolar spindles in *Ccnb3^−/−^* oocytes retained the barrel-shaped appearance characteristic of metaphase I even at late time points 16 hours after GVBD, when control oocytes had progressed to metaphase II (**Figure 2C**).

### Cell cycle arrest in Ccnb3^−/−^ oocytes is due to incomplete APC/C activation

Next, we wished to determine the molecular basis of the failure in chromosome segregation in *Ccnb3^−/−^*oocytes. Separase is a cysteine protease that cleaves the kleisin subunit of cohesin complexes to dissolve sister chromatid cohesion at anaphase onset in mitosis and meiosis [29]. Prior to anaphase, separase activity is restrained in part by binding to its inhibitory chaperone, securin, and by high cyclin B-CDK activity [29]. In oocytes, the APC/C-dependent degradation of securin and cyclin B1 leads to activation of separase, which then cleaves the meiotic kleisin REC8, allowing the separation of homologous chromosomes [30–32]. To follow separase activity in live oocytes, we expressed a separase activity sensor we recently developed[33], similar to one described in mitotic tissue culture cells [34]. The sensor harbors two fluorescent tags separated by the mitotic cohesin subunit RAD21 as a separase cleavage substrate; this ensemble is fused to histone H2B to allow it to localize to chromosomes. Upon cleavage by separase, colocalization of the fluorophores is lost, leaving only the H2B-attached fluorophore on chromosomes (**Figure 3A**). Sensor cleavage occurs only under conditions when endogenous REC8 is removed from chromosome arms[33]. When control oocytes in culture were injected with mRNA encoding the sensor, the fluorescent signal on chromosomes changed abruptly from yellow to red upon anaphase onset (**Figure 3B**, top row, compare 6:20 to 6:40 time points). In contrast, both fluorophores of the sensor remained chromatin-associated throughout the arrest in *Ccnb3^−/−^* oocytes (**Figure 3B**, bottom row), indicating that separase cannot be activated in meiosis I in the absence of cyclin B3, explaining in turn why the mutant oocytes cannot separate homologous chromosomes and extrude PBs.

**Figure 3.**
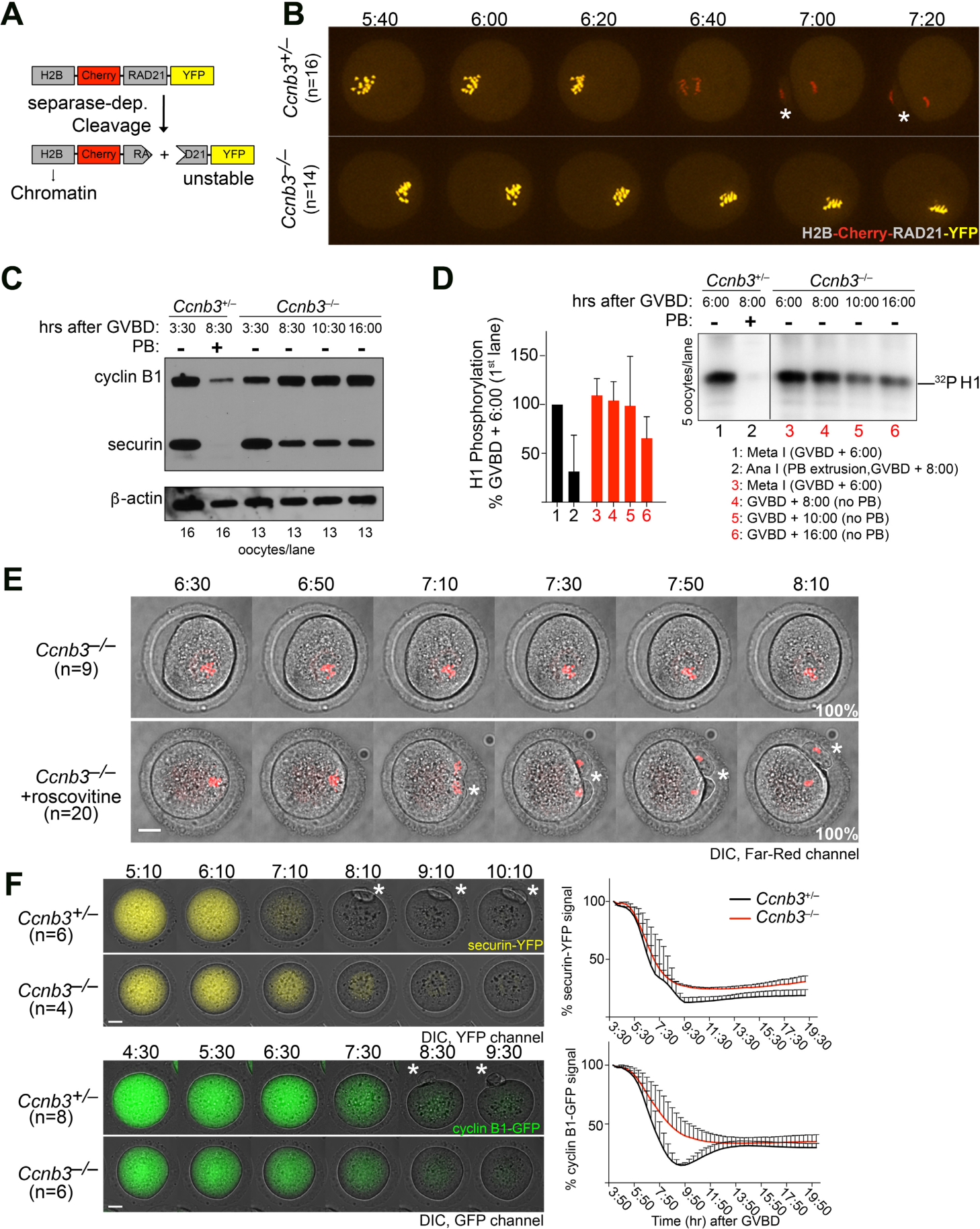
Cell cycle arrest in *Ccnb3 ^−/−^* oocytes is due to incomplete APC/C activation. (A) Schematic of the separase activity sensor. See text for details. (B) Failure to activate separase in the absence of cyclin B3. Separase activity sensor mRNA was injected into GV oocytes, which were released into meiosis I and visualized by spinning disk confocal microscopy. Selected time frames of collapsed z-sections (11 section, 3-µm steps) from a representative movie are shown. Time points after GVBD are indicated as hours:minutes. Scale bar represents 20 µm; white asterisks indicate PBs. (C) Western blot analysis of cyclin B1 and securin during oocyte maturation, at the time points indicated as hours:minutes after GVBD. β-actin serves as loading control. The number of oocytes used and the presence or absence of a PB are indicated. (D) Total MPF activity during oocyte maturation, at the time points indicated as hours:minutes after GVBD. Histone H1 was used as a substrate. Five oocytes were used per kinase reaction, and the presence or absence of a PB is indicated. The graph on the side shows the quantification of phosphate incorporation from 3 independent experiments and error bars indicate standard deviation. (E) CDK inhibition rescues meiosis I division. Oocytes were incubated with SiR-DNA to visualize chromosomes. In metaphase I, 6 hours 20 minutes after GVBD, oocytes were treated with 0.2 mM roscovitine (final concentration), where indicated, and the movie was started. Selected time frames of collapsed z-sections (11 section, 3-µm steps) of the DIC and far-red channel from a representative movie of *Ccnb3^−/−^* oocytes with or without roscovitine treatment are shown. The asterisk indicates chromosome segregation in anaphase I. Time points after GVBD are indicated as hours:minutes. Scale bar represents 20 µm. (F) Degradation of exogenous APC/C substrates. Securin-YFP (above) or cyclin B1-GFP (below) mRNA were injected into GV oocytes. Stills from representative movies are shown. Time points after GVBD are indicated as hours: minutes. Scale bar represents 20 µm (n: number of oocytes) and white asterisk indicates polar body extrusion (PBE). Quantification of fluorescence intensities is shown on the right (mean ± s.d. from the indicated number of oocytes imaged).

The failure to activate separase was potentially due to a failure to degrade cyclin B1 and securin, the two inhibitors of separase. To test this, we analyzed levels of each protein by western blotting (**Figure 3C**). In control oocytes, the levels of both proteins decreased precipitously at metaphase-to-anaphase onset (compare 8:30 to 3:30 after GVBD). In *Ccnb3^−/−^* oocytes in contrast, cyclin B1 levels did not decrease at any point during culturing, even at late time points. In fact, cyclin B1 levels were consistently higher during arrest (8:30 and later) as compared to prometaphase (3:30). Securin levels decreased between 3:30 and 8:30 after GVBD, but remained at much higher levels than in the controls. Thus, although securin levels drop partially, both proteins are inappropriately stabilized in cyclin B3-deficient cells at the time wild-type cells would normally carry out the meiosis I division.

Next we asked whether stabilization of cyclin B1 in *Ccnb3^−/−^* oocytes correlated with elevated kinase activity of MPF (M-phase promoting factor), a heterodimer of CDK1 and either cyclin B1 or B2 (usually called simply "cyclin B") [1, 4]. For this we performed *in vitro* kinase assays using oocyte extracts at the indicated time points to assess MPF activity [35] (**Figure 3D**). As previously shown, control oocytes showed high MPF activity in metaphase I (6:00 after GVBD), and a drop in kinase activity when they extrude PBs (8:00 after GVBD) [36]. *Ccnb3^−/−^* oocytes similarly showed high MPF activity in metaphase I (6:00 after GVBD), but no drop in kinase activity was observed at the time when normal cells extruded PBs (8:00 after GVBD) (**Figure 3D**).

Because endogenous securin levels decreased slightly in *Ccnb3^−/−^* oocytes (**Figure 3C**), whereas cyclin B1 levels and MPF activity did not (**Figure 3C,D**) at a time when control oocytes extrude PBs, we asked whether inhibiting MPF activity at the normal time of anaphase I onset would be sufficient for *Ccnb3^−/−^* oocytes to undergo the metaphase I-to-anaphase I transition and PB extrusion. Indeed, addition of the CDK inhibitor roscovitine [37] rescued PB extrusion and chromosome segregation in *Ccnb3^−/−^* oocytes, suggesting that the failure to undergo anaphase I without cyclin B3 is mainly attributable to persistently elevated MPF activity (**Figure 3E**).

Collectively, the results thus far lead us to conclude that *Ccnb3^−/−^* oocytes that do not extrude PBs are arrested in metaphase I with aligned homologous chromosome pairs, bipolar metaphase I spindles, high securin and cyclin B1 protein levels, and little or no separase activity. In oocytes, degradation of cyclin B1 and securin depends on their ubiquitination by the APC/C [31, 32]. Hence, our results suggested that without cyclin B3, the APC/C is not sufficiently activated on time to properly target endogenous cyclin B1 and securin. Because endogenous securin levels did decrease slightly in the mutant, however, we suspected that the APC/C might be partially active. To test this, we asked whether exogenously expressed securin and cyclin B1 were also stabilized in *Ccnb3^−/−^* oocytes that were arrested in metaphase I. mRNAs coding for either protein fused to fluorescent tags were injected into GV oocytes and followed by live imaging [30, 32, 38–41]. Surprisingly, cyclin B1-GFP and securin-YFP were both efficiently degraded in *Ccnb3^−/−^* oocytes (**Figure 3F**). Securin-YFP was degraded with kinetics and final extent similar to the controls, whereas cyclin B1-GFP appeared to be degraded slightly more slowly and was higher in *Ccnb3^−/−^* than in control cells specifically at the time the controls underwent PB extrusion (**Figure 3F**). Thus, even though both exogenous substrates were more efficiently degraded than their endogenous counterparts, the differences between the two substrates were similar in both contexts (i.e., securin was degraded to a greater extent than cyclin B1 in the absence of cyclin B3). These results indicate that the APC/C does indeed become partially active in the mutants, but the data further suggest that cyclin B3 in some manner specifically promotes the degradation of endogenous APC/C substrates. This may be by modulating APC/C specificity per se, or by regulating the substrates or their localization to affect their ability to be ubiquitylated. This role of cyclin B3 appears to be largely dispensable in the case of exogenously expressed, tagged substrates.

Timely APC/C activation depends on satisfaction of the SAC [28, 42], so we asked whether inappropriate SAC activity was the reason for arrest in *Ccnb3^−/−^* oocytes. Kinetochore recruitment of SAC components, such as MAD2, is a readout for SAC activation [43]. We found that MAD2 was recruited to unattached kinetochores early in meiosis I [40, 44] in both control and *Ccnb3^−/−^* oocytes, and that neither control nor *Ccnb3^−/−^*oocytes retained MAD2 at kinetochores in metaphase I (**Figure 4**). We conclude that *Ccnb3^−/−^* oocytes have a functional SAC (i.e., it can recognize unattached kinetochores in early meiosis I) that is appropriately satisfied upon kinetochore-microtubule attachment at metaphase I. Thus, prolonged SAC activation is not the reason for the metaphase I arrest and the failure to efficiently degrade endogenous APC/C substrates in *Ccnb3^−/−^* oocytes.

**Figure 4.**
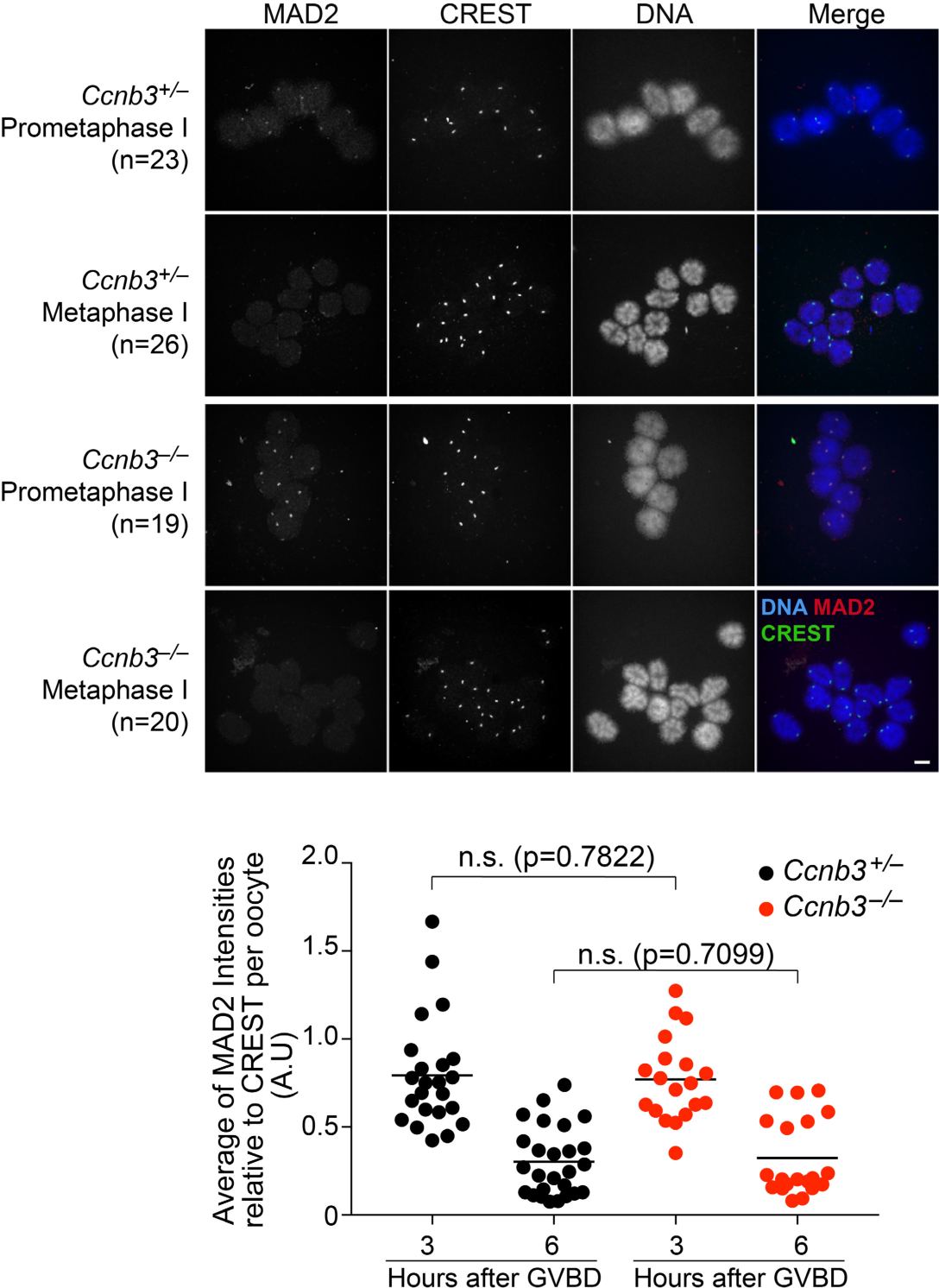
SAC activation is not the cause of metaphase I arrest in *Ccnb3 ^−/−^* oocytes. (Top) Chromosome spreads were prepared 3 hours (early prometaphase I) and 6 hours (metaphase I) after GVBD, then stained with Hoechst (blue), CREST (green), and anti-MAD2 (red). Scale bar represents 5 µm. (Bottom) Quantification of MAD2 signal intensity relative to CREST; each point is the mean relative intensity averaged across centromeres in an oocyte. P values are from t tests (n.s., not significant).

### Cyclin B3 is a later APC/C substrate than securin and cyclin A2

Like other M-phase cyclins, cyclin B3 harbors a conserved destruction box (D-box) [18], a motif typically targeted for APC/C-dependent ubiquitination [14] (**Figure 1A**). In mitosis, exogenously expressed cyclin B3 was shown to be degraded after cyclin B1[18]. We therefore asked whether cyclin B3 protein stability is regulated in a cell cycle-dependent manner in oocytes. mRNA encoding cyclin B3 tagged with RFP was injected into wild-type oocytes together with mRNA for either securin-YFP or cyclin A2-GFP, and oocytes were followed by live imaging. Exogenously expressed cyclin B3-RFP accumulated during meiosis I, then was degraded abruptly just before PB extrusion. Importantly, cyclin B3 degradation occurred after degradation of both cyclin A2 and securin (**Figure 5A and B**), showing temporally ordered degradation of APC/C substrates in oocyte meiosis I. No re-accumulation of cyclin B3-RFP was observed as oocytes progressed into meiosis II. Deleting the D-box stabilized cyclin B3-RFP (**Figure 5C**) (the protein has no obvious KEN-box or ABBA motif [45]. These findings support the conclusion that cyclin B3 is an APC/C substrate in oocyte meiosis.

**Figure 5.**
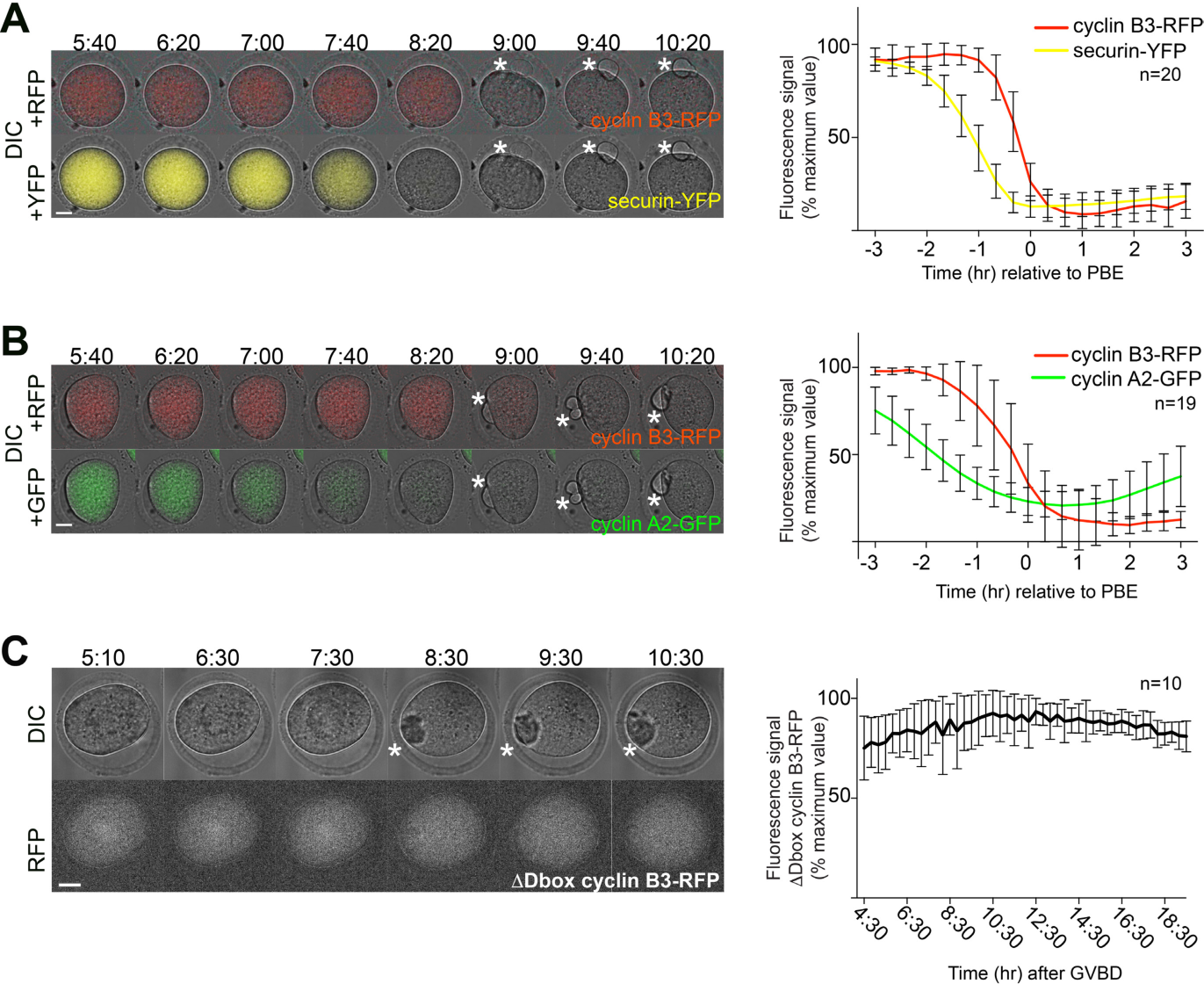
Ordered degradation of APC/C substrates in oocyte meiosis. I. (A,B) Cyclin B3 is degraded after securin or cyclin A2. mRNA encoding cyclin B3-RFP and either securin-YFP (panel A) or cyclin A2-GFP (panel B) were co-injected into GV oocytes of CD-1 mice. Each panel shows selected frames from a movie of a representative oocyte, with an overlay of DIC and RFP channels in the top rows, and DIC and YFP or GFP channels in the bottom rows. (C) The D box of cyclin B3 is required for degradation. CD-1 GV oocytes were injected with mRNA encoding cyclin B3-RFP with the D box deleted (ΔDbox). Frames from a representative movie are shown, with the DIC channel on top and RFP at the bottom. All panels: Time points after GVBD (BD) are indicated as hours:minutes.Scale bar represents 20 μm and white asterisks indicate PBs. Quantification of fluorescence intensities is shown on the right (mean ± s.d. of the indicated number of oocytes).

### Cyclin B3-associated kinase activity is required for progression through oocyte meiosis I

To determine whether mouse cyclin B3 can support CDK1-dependent kinase activity (as for the *C. elegans*, *Drosophila*, and chicken proteins [13, 17, 46]), we co-expressed and purified recombinant murine protein complexes from insect cells. We expressed cyclin B3 that was N-terminally tagged with maltose binding protein (MBP) and hexahistidine (^MBPHis^cyclin B3) and untagged CDK1 or a hemagglutinin (HA) tagged version of CDK1 that runs slightly slower than untagged on SDS-PAGE. Both CDK1 and CDK1-HA could be co-purified with ^MBPHis^cyclin B3 upon affinity purification on amylose resin (**Figure 6A**, left, lanes 2-3), and both types of cyclin B3-CDK1 complex displayed kinase activity towards histone H1 *in vitro* (**Figure 6A**, right, lanes 2-3). To further investigate this kinase activity, we mutated the MRAIL motif on cyclin B3, which is conserved among different cyclins and is located on the hydrophobic patch formed by the cyclin fold [47, 48]. MRAIL mutations were previously shown to prevent binding of cyclin-CDK complexes to potential substrates without affecting cyclin-CDK interactions [47], although other studies also reported loss of cyclin-CDK interaction [49][61]. Mutating the methionine, arginine, and leucine residues of the MRAIL motif to alanine (cyclin B3-MRL mutant) did not affect the ability to co-purify CDK1 or CKD1-HA (**Figure 6A**, left, lanes 5-6), but the affinity-purified ^MBPHis^cyclin B3 MRL-CDK complexes were not active in the histone H1 kinase assay (**Figure 6A**, right, lanes 5-6). Furthermore, affinity purification of wild-type cyclin B3 without co-expressed CDK1 yielded detectable H1 kinase activity, presumably attributable to cyclin B3 association with an endogenous insect cell CDK(s), but the MRL mutant gave no such baseline activity (**Figure 6A**, right, lanes 1 and 4).

**Figure 6.**
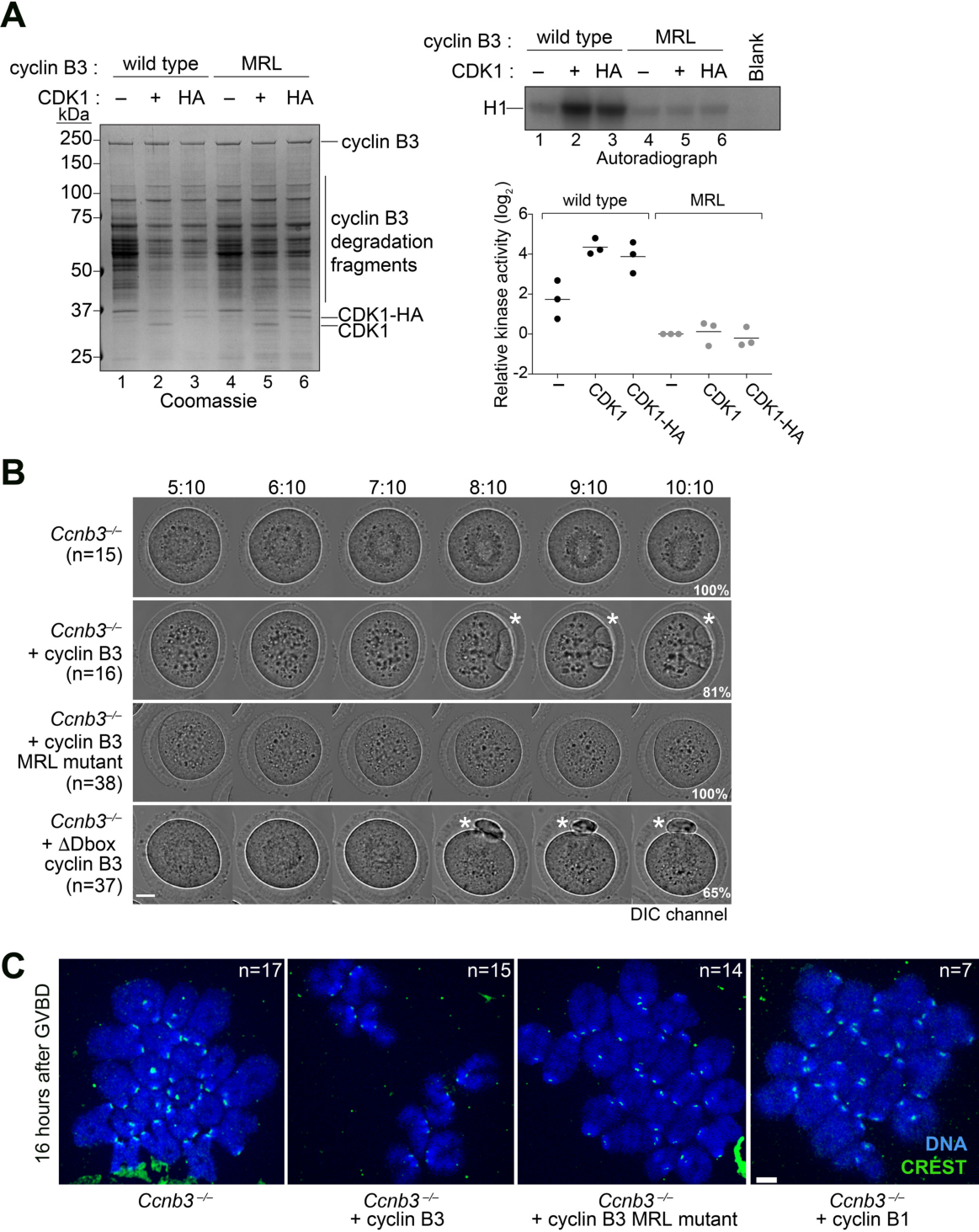
Only cyclin B3 that can support *in vitro* kinase activity can rescue Ccnb3 ^−/−^ oocytes. (A) Affinity purification of cyclin B3-CDK1 complexes. ^MBPHis^cyclin B3 or ^MBPHis^cyclin B3 MRL mutant were expressed in insect cells alone or co-expressed with either untagged or HA-tagged CDK1. (Left) The eluates from purification on amylose resin were separated on SDS-PAGE and stained with Coomasssie. (Right) Representative autoradiograph (top) and quantification (bottom) from histone H1 kinase assays. In the graph, values in each experiment (n =3) were normalized to the signal from the ^MBPHis^cyclin B3 MRL sample (lane 4 in the autoradiograph). (B, C) MRL-mutant cyclin B3 fails to rescue Ccnb3^−/−^ oocytes. Ccnb3^−/−^ oocytes were sham-injected or injected with the indicated cyclin mRNA, then released into meiosis I. Frames of representative movies are shown in panel B. Times after GVBD are indicated as hours:minutes and percentages of oocytes of the shown phenotypes are indicated. Scale bar represents 20 μm and white asterisks indicate PBs. Panel C shows representative chromosome spreads 16 hours after GVBD. In the case of oocytes injected with the wild type cyclin B3 construct, only oocytes that had extruded PBs were analyzed. Chromosomes were stained with Hoechst (blue) and kinetochores with CREST (green). Scale bar represents 5μm.

We conclude that cyclin B3 can form an active kinase complex with CDK1, suggesting in turn that CDK1 may be a physiological partner of cyclin B3 *in vivo*. We also conclude that the MRL mutant permits CDK1 binding but abolishes associated kinase activity.

We further tested whether oocyte division requires cyclin B3-associated kinase activity. To this end, we first performed live imaging on *Ccnb3^−/−^* oocytes injected with mRNA encoding wild-type cyclin B3 or protein with the D-box deleted (ΔD-box). Both constructs efficiently rescued PB extrusion in the mutant (**Figure 6B**). Chromosome spreads demonstrated that separation of homologous chromosomes was also rescued upon cyclin B3 expression (**Figure 6C**). This finding confirmed that meiotic arrest is attributable to absence of cyclin B3 *per se* and established feasibility of structure/function experiments based on complementation of *Ccnb3^−/−^* oocyte arrest. In striking contrast to wild-type cyclin B3, injection of mRNA encoding the MRL mutant was unable to rescue meiosis I division, PB extrusion, or chromosome segregation (**Figure 6B and C**), indicating that cyclin B3-CDK complexes (likely CDK1) bring about anaphase I onset through their kinase activity towards yet unknown substrates. Note that, unlike cyclin B3, expression of exogenous cyclin B1 was unable to rescue PB extrusion (**Figure 3F**) or homolog disjunction (**Figure 6C**). We infer that cyclin B3-CDK1 substrates are likely distinct from cyclin B1-CDK1 substrates, perhaps due to distinct substrate specificity or accessibility.

### Functional conservation of vertebrate cyclin B3 in meiosis I

It was previously reported that *X. laevis* oocytes contain cyclin B3 mRNA but not detectable levels of the protein, from which it was inferred that cyclin B3 plays no role in Xenopus oocyte meiosis [12]. Furthermore, mammalian cyclin B3 is much larger than cyclin B3 from non-mammalian vertebrates due to the presence of the extended exon 7 [10], suggesting that the protein might be functionally distinct in mammals. To explore this possibility, we tested for inter-species cross-complementation by injecting mRNAs encoding cyclin B3 from either *X. laevis* or zebrafish (*Danio rerio*) into mouse *Ccnb3^−/−^* oocytes. Remarkably, heterologous expression of cyclin B3 from either species efficiently rescued PB extrusion (**Figure 7A)**. Live imaging of oocytes stained with SiR-DNA to visualize chromosomes showed no obvious defect in chromosome alignment or anaphase I onset in oocytes rescued with *X. laevis* cyclin B3 (**Figure 7B**). These findings demonstrate that the extended exon 7 in mouse cyclin B3 is dispensable to promote anaphase I onset. We further infer that biochemical functions of cyclin B3 protein are conserved across vertebrates, in turn raising the possibility that cyclin B3 promotes the oocyte meiosis I division throughout the vertebrate lineage. Even more remarkably, Drosophila cyclin B3 was also able to rescue the mouse mutant oocytes (**Figure 7A**), suggesting conservation of biochemical properties throughout metazoa.

**Figure 7.**
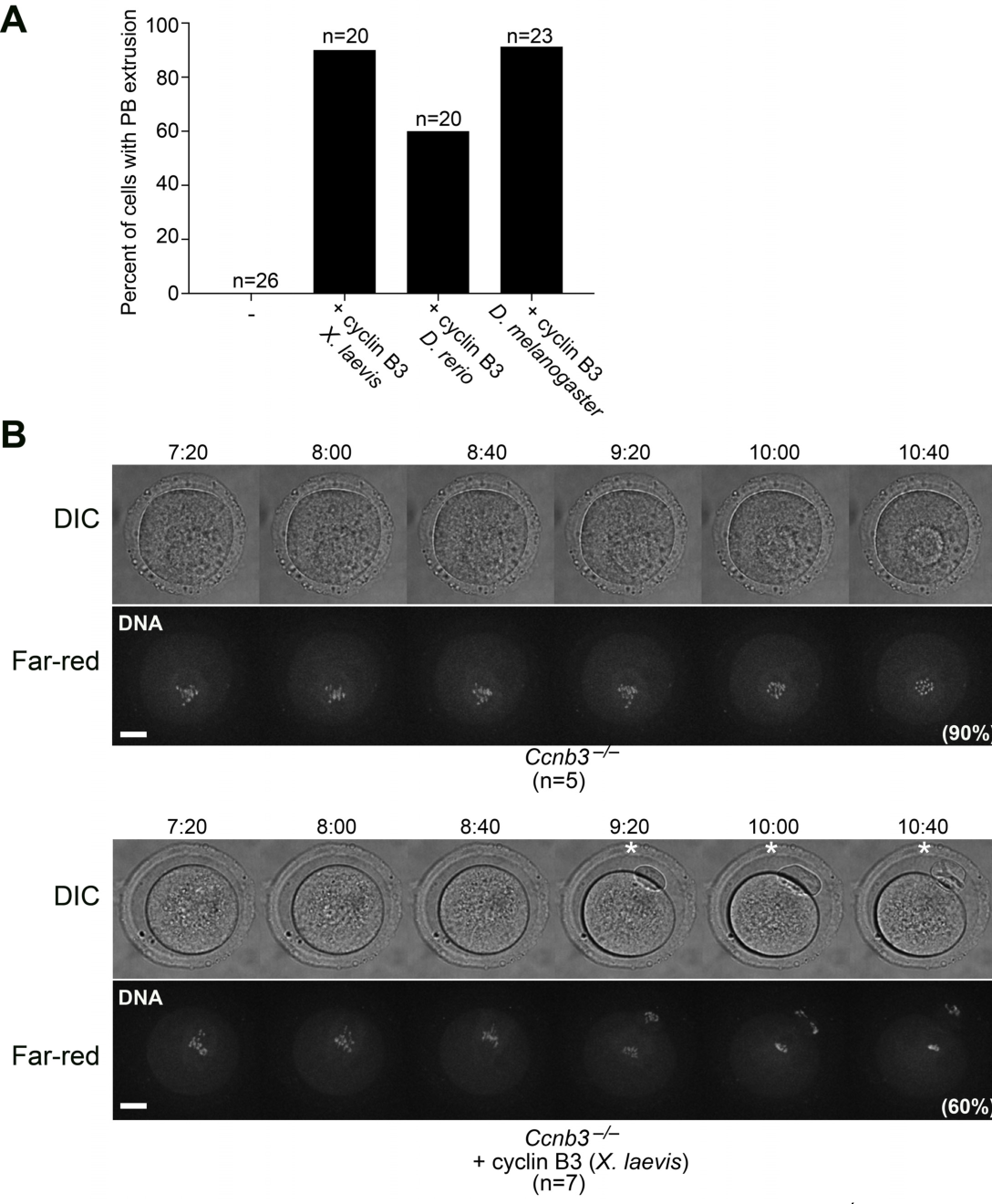
Inter-species cross-complementation of *Ccnb3 ^−/−^* oocytes. (A) *Ccnb3^−/−^* oocytes were injected with the indicated mRNA, induced to enter meiosis I, and scored for PB extrusion. (**B**) Selected time frames of collapsed z-sections (12 sections, 3-µm steps) from a representative spinning disk confocal movie of *Ccnb3^−/−^* sham-injected oocytes, and *Ccnb3^−/−^* oocytes injected with *X. laevis* cyclin B3 mRNA. Prior to live imaging oocytes were incubated with SiR-DNA. Top panel shows the DIC channel and bottom panel shows SiR-DNA staining in far-red. Time points after GVBD are indicated as hours:minutes. Scale bar represents 20 µm and white asterisks indicate PBs.

## Discussion

The aim of our study was to determine whether cyclin B3 is specifically required for meiosis, using a knock-out strategy to create mice devoid of the protein. Whereas loss of cyclin B3 has no effect on male fertility (Karasu, 2018, in preparation), mutant females are sterile because cyclin B3 promotes APC/C activation and thus anaphase onset in the first meiotic division in oocytes. Similar findings were obtained independently with a different targeted mutation in *Ccnb3* (see accompanying manuscript by Li et al.). Our findings definitively establish a crucial role for mouse cyclin B3 in the female germline.

The requirement in anaphase I onset in mouse is similar to *Drosophila* oocytes [14, 16]. It has been proposed that cyclin B3 regulates APC/C activity during the rapid mitotic cycles of the early *Drosophila* embryo [14]. Our data are in accordance with a similar role in mouse oocyte meiosis I. Interestingly, however, cyclin B3-deficient oocytes show a distinct partial APC/C activation phenotype in which exogenous APC/C targets are degraded efficiently but endogenous substrates are not. To our knowledge this is the first time that such a discrepancy has been observed in mouse oocytes. It is not yet clear what distinguishes exogenous from endogenous substrates. One possibility is that expression timing (i.e., starting upon mRNA injection into GV oocytes) places the exogenous proteins at a disadvantage relative to endogenous proteins for binding to essential partners, e.g., of securin to separase and cyclin B1 to CDK. It is also possible that the fluorescent protein tags interfere slightly with this binding. It is noteworthy in this context that free securin is degraded earlier than separase-bound securin, which is stabilized by maintenance in a non-phosphorylated state through association with phosphatase PP2A [50]. Securin phosphorylation enhances its affinity for the APC/C [50, 51]. Regardless of the source of the distinction, our data clearly demonstrate that APC/C becomes partially active on time, but not sufficiently active to robustly provoke degradation of all of its intended targets and to trigger anaphase I.

Why is fine tuning of APC/C activity by cyclin B3 important in oocyte meiosis? APC/C in association with CDH1 is required to maintain oocyte arrest in prophase I by destabilizing cyclin B1 and Cdc25 [52–54]. At meiosis I entry, CDH1 is only gradually shut off, leading to a slow increase of MPF (cyclin B1-CDK1) kinase activity [52, 55–57], and not the sharp rise observed in mitotic cells. Once MPF activity is high enough, CDH1 is inactivated. At this time in prometaphase I, SAC control keeps APC/C-CDC20 in check to prevent precocious securin and cyclin B degradation [41, 58–60]. In mice, the slow handover from APC/C-CDH1-dependent to APC/C-CDC20-dependent ubiquitination is specific to oocytes. Furthermore, unlike in mitosis, SAC control is leaky in oocyte meiosis I, and onset of APC/C-dependent degradation occurs before full kinetochore attachment status has been achieved [42]. Because of these specificities, cyclin B3 may be required in oocytes, and only in oocytes, to ensure the sudden and full activation of separase by mediating rapid and efficient degradation of the pool of securin and cyclin B1 that is inhibiting separase after the degradation of free securin and cyclin B1 in the cytoplasm.

We propose that cyclin B3 is required to tip the balance towards synchronized, full APC/C activity and progression into anaphase. Without cyclin B3 the APC/C is active, but endogenous cyclin B1 is not at all, and securin not efficiently, targeted for degradation. Consistent with this view, the rescue of *Ccnb3^−/−^* oocytes by artificially downregulating MPF activity with roscovitine indicates that sufficient separase has been liberated from its inhibitory interaction with securin in the mutant. Cyclin B3 in mammalian oocytes appears dispensable for turning off the SAC, unlike in *C. elegans* [16], so we consider it more likely that cyclin B3-CDK1 complexes foster full APC/C activity directly by specifically modifying APC/C substrates, APC/C subunits, APC/C activators (CDC20 and CDH1), or some combination [61]. Nevertheless, cyclin B3 is not strictly essential, in that a small proportion of mice harbored oocytes that were able to progress through meiosis I. The reason for this incomplete penetrance is not yet clear, but may reflect subtle variation (perhaps from strain background) in either the degree of APC/C activation or the threshold of APC/C activity needed for progression. Put another way, we suggest that cyclin B3-deficient oocytes occasionally achieve just enough APC/C activity during a critical time window to cross a threshold needed for the switch-like transition into anaphase I.

In budding and fission yeast, sequential substrate phosphorylation during cell cycle progression is thought to be brought about primarily by changes in overall CDK activity levels as opposed to different substrate specificities of different cyclins [2, 62–64]. Accordingly, the whole cell cycle can be driven by a single CDK-cyclin fusion complex in *S. pombe* [62, 65]. In mouse it was shown that CDK1 is the only CDK essential for cell cycle progression *per se,* in mitosis [66]. Loss of cyclin B2 does not affect viability or fertility, showing that cyclin B1 can substitute [67]. It may thus seem surprising that cyclin B3 should have such a specific role during oocyte meiosis that cannot be replaced by cyclin B1. We can rule out the possibility that exogenous cyclin B1 failed to rescue *Ccnb3^−/−^* oocytes simply because cyclin B1 is normally degraded earlier than cyclin B3: the persistence of high levels of endogenous cyclin B1 in the *Ccnb3^−/−^* oocytes showed that cyclin B1 was indeed present in the rescue experiment and thus should have restored anaphase onset if cyclin B1 were capable of substituting for cyclin B3. Hence, cell cycle progression in meiosis appears to be specifically dependent on individual CDK complexes. This conclusion is further supported by the fact that loss of CDK2 does not affect somatic cells but leads to sterility due to failure to enter meiosis I [68]. Cell type specificity is also seen for cyclin A2, which is required in hematopoetic stem cells, fibroblasts, and female meiosis, but not in other tissues [21, 69, 70].

Overall, our data show that cyclin B3 is an M-phase cyclin with a singular role in oocyte meiosis, probably due to the specificities of meiotic cell cycle regulation in this huge cell. A key challenge now will be defining which set of substrates needs to be phosphorylated by cyclin B3-CDK complexes to bring about proper meiotic progression.

## Materials and Methods

### Animals

Mice in the SK lab were maintained and sacrificed for the experiments under regulatory standards approved by the Memorial Sloan Kettering Cancer Center (MSKCC) Institutional Animal Care and Use Committee. Mice in the KW lab were maintained according to current French guidelines in a conventional mouse facility (UMR7622), with the authorization C 75-05-13. All experiments were subject to ethical review and approved by the French Ministry of Higher Education and Research (authorization n° B-75-1308). For this study females were either purchased at 7 weeks of age (CD-1 mice, Janvier Lab France), and used at 8-16 weeks of age for experiments, or bred in our animal facilities and used at 8-16 weeks of age (*Ccnb3* mouse strain). Except for genotyping, the mice had not been involved in any previous procedures prior to experimentation, they were given *ad libitum* access to food and water supply. Housing was done under a 12 hour light / 12 hour dark cycle according to the Federation of European Laboratory Science Associations (FELASA).

### Generation of Ccnb3 knock-out mouse strain, husbandry and genotyping

Endonuclease mediated (*em*) allele was generated by the MSKCC Mouse Genetics Core Facility. Exon 7 (Exon numbering used here is from the current accession. The targeted exon corresponds to exon 8 in the numbering used by Lozano et al. [10] of *Ccnb3* (NCBI Gene ID: 209091) was targeted by guide RNA (C40-TGAACTTGGCATGATAGCAC). Guide RNA was cloned into pDR274 vector for in vitro transcription. In vitro transcribed guide RNA (10 ng/μl) and Cas9 (20 ng/μl) were microinjected into pronuclei of CBA/J × C57BL/6J F2 hybrid zygotes using conventional techniques [74].

Genomic DNA from founder animals were subjected to PCR by using following primers, CCN3B-A GAGTATTAGCACTGAGTCAGGGAC and CCN3B-B GGAATACCTCAGTTTCTTTTGCAC and T7 endonuclease I digestion was performed to identify the animals carrying indels. Since male animals have one copy of the X-linked *Ccnb3* gene, T7 assay on males was performed in the presence of wild type genomic DNA.

To define the molecular nature of indels, genomic DNA from T7-positive animals was amplified using the primers indicated above. Amplified PCR fragments were used for TA Cloning (TA Clonining™ Kit with pCR™ 2.1 vector, Invitrogen). Ten white bacterial colonies were selected, inserts were sequenced, and the mutations were characterized as deletions, insertions or substitutions. The *em1* allele, an out of frame deletion, was chosen to generate the *Ccnb3^em1Sky^* line. After two rounds of backcrossing to C57BL/6J, animals were interbred to generate homozygous and heterozygous female animals and hemizygous male animals. Primers for genotyping were GT4-F TGTTGATGAAGAGGAATTTTTCAAATCATTCCT and GT4-R TTCTTTTGCACCCAGAGTTGACTTAAAG. The amplified PCR product was subjected to BsrI enzyme digestion (NEB). A BsrI site is lost in the *em1* allele. Since *Ccnb3* is X-linked, no *Ccnb3^+/+^*females can be obtained through crosses yielding homozygous *Ccnb3^−/−^* mice. Therefore, we bred *Ccnb3^−/Y^* males with *Ccnb3^+/–^* females to obtain *Ccnb3^−/−^* and *Ccnb3^+/–^* females to use as experimental and littermate mate controls, respectively.

### Oocyte harvesting and in vitro culture

GV stage oocytes were collected from ovaries of sacrificed CD-1 mice or *Ccnb3^+/–^* and *Ccnb3^−/−^* mice aged 8-16 weeks. Ovaries were transferred to homemade M2 medium at a temperature of 38°C and dissected to obtain oocytes. Oocytes were collected and isolated from follicular cells with mouth pipetting using torn-out Pasteur pipettes. For all microinjection experiments, ovaries and oocytes were incubated in commercial M2 medium (Sigma-Aldrich) and kept at GV stage by the addition of dibutyl cyclic AMP (dbcAMP, Sigma-Aldrich) at 100 µg/ml final concentration. Oocytes undergoing GVBD within at most 90 min after ovary dissection or release from dbcAMP were used for experiments. For experiments using *Ccnb3^−/−^* mice, 5 oocytes per mouse were left untreated and were released into meiosis I to ascertain that no PB extrusion occurred. Roscovitine was added 6h20 after GVDB at a final concentration of 0,2 mM.

### Histology

Ovaries from *Ccnb3^+/–^* and *Ccnb3^−/−^* adult mice were fixed overnight in 4% paraformaldehyde (PFA) at 4°C. Fixed tissues were washed in water and stored in 70% ethanol at 4°C. Fixed tissues were submitted to the Molecular Cytology Core Facility (MSKCC) for embedding in paraffin and sectioning of the embedded tissue (whole ovary) as 8 µm sections and preparation of slides with ovary sections. The slides were then subjected to immunohistochemistry as follows: The immunohistochemical staining was performed at Molecular Cytology Core Facility (MSKCC) using a Discovery XT processor (Ventana Medical Systems). Ovary sections were de-paraffinized with EZPrep buffer (Ventana Medical Systems), antigen retrieval was performed with CC1 buffer (Ventana Medical Systems). Sections were blocked for 30 minutes with Background Buster solution (Innovex), followed by avidin-biotin blocking for 8 minutes (Ventana Medical Systems). Sections were then incubated with anti-VASA (Abcam, cat#ab13840, 0.17 µg/ml) for 5 hours, followed by 60 minutes incubation with biotinylated goat anti-rabbit (Vector Labs, cat# PK6101) at 1:200 dilution. The detection was performed with DAB detection kit (Ventana Medical Systems) according to manufacturer instruction. Slides were counterstained with hematoxylin and cover slipped with Permount (Fisher Scientific). IHC slides were scanned using Pannoramic Flash 250 with a 20×/0.8 NA air objective. Images were analyzed using Pannoramic Viewer Software (3DHistech). To count oocytes, every fifth section on the slide was analyzed and the number of oocytes in primordial, primary, secondary and antral follicles were noted and summed.

### Histological examination of somatic tissues

Gross histopathological analysis of major organs and tissues was performed by the MSKCC Laboratory of Comparative Pathology Core Facility for the following male mice: two 2-month old *Ccnb3^−/Y^* males and one 2-month old *Ccnb3^+/Y^* male. Histological examination of following tissues was perfomed: diaphragm, skeletal muscle, sciatic nerve, heart/aorta, thymus, lung, kidneys, salivary gland, mesenteric lymph nodes, stomach, duodenum, pancreas, jejunum, ileum, cecum, colon, adrenals, liver, gallbladder, spleen, uterus, ovaries, cervix, urinary bladder, skin of dorsum and subjacent brown fat, skin of ventrum and adjacent mammary gland, thyroid, parathyroid, esophagus, trachea, stifle, sternum, coronal sections of head/brain, vertebrae and spinal cord. Tissues were fixed in 10% neutral buffered formalin and bones were decalcified in formic acid solution using the Surgipath Decalcifier I (Leica Biosystems) for 48 hr. Samples were routinely processed in alcohol and xylene, embedded in paraffin, sectioned (5 mm), and stained with hematoxylin and eosin. Examined tissue and organs were grossly normal in both *Ccnb3^−/Y^* males and *Ccnb3^+/Y^* male.

### Extract preparation and western blotting

To analyze endogenous protein levels, oocytes were cultured as described above. Oocytes were harvested at the indicated time points, washed in phosphate buffer saline (PBS) to remove proteins from the medium and then placed on the wall of the Eppendorf tube and snap-frozen in liquid nitrogen. Later, 7.5 µl 1× Laemmli lysis buffer was added to the tube and samples were boiled 5 minutes at 100°C.

For western blotting, samples were separated on 4–12% Bis-Tris NuPAGE precast gels (Life Technologies) at 150 V for 70 minutes. Proteins were transferred to polyvinylidene difluoride (PVDF) membranes by wet transfer method in Tris-Glycine-20% methanol, at 120 V for 40 minutes at 4°C. Membranes were blocked with 5% non-fat milk in PBS-0.1% Tween (PBS-T) for 30 minutes at room temperature on an orbital shaker. Blocked membranes were incubated with primary antibodies 1 hr at room temperature or overnight at 4°C. Membranes were washed with PBS-T for 30 minutes at room temperature, then incubated with HRP-conjugated secondary antibodies for 1 hr at room temperature. Membranes were washed with PBS-T for 15 minutes and the signal was developed by ECL Plus Perkin Elmer or ECL Prime GE Healthcare.

To analyse protein levels from mouse testis extracts, dissected testes were placed in an Eppendorf tube, frozen on dry ice and stored at −80°C. The frozen tissue was resuspended in RIPA buffer (50 mM Tris-HCL pH 7.5, 150 mM NaCl, 0.1% SDS, 0.5% sodium deoxycholate, 1% NP40) supplemented with protease inhibitors (Roche, Mini tablets). The tissue was disrupted with a plastic pestle and incubated by end-over-end rotation for 15 minutes, at 4°C. After brief sonication, samples were centrifuged at 15000 rpm for 15 minutes. The clear lysate was transferred to a new tube, and used for immunoprecipitation. Whole cell extract was pre-cleared with protein G beads by end-over-end rotation for 1 hr, at 4°C. Antibodies were added to pre-cleared lysates and incubated overnight with end-over-end rotation, at 4°C. Protein G beads were added to the tubes and incubate for 1 hr with end-over-end rotation, at 4°C. Beads were washed three times with RIPA buffer, resuspended in 1x NuPAGE LDS sample buffer (Invitrogen) with 50 mM DTT, and boiled 5 minutes to elute immunoprecipitated proteins.

Primary antibodies were used at the following dilutions to detect proteins: mouse anti-securin (ab3305, Abcam, 1:300), mouse anti-cyclin B1 (ab72, Abcam, 1:400), mouse anti-β actin (8H10D10, CST,1:1000), mouse-anti-cyclin B3 (Abmart Inc, 1:500), and rat anti-tubulin alpha (MCA78G, Biorad, 1:1000). Secondary antibodies were used at the following dilutions: goat anti-mouse IgG (H+L)-HRP (1721011, Biorad, 1:10000), and goat anti-rat IgG-HRP (AP136P, Millipore, 1:10000).

### Plasmids

Mouse *Ccnb3* mRNA was amplified by PCR from whole testis cDNA library and cloned into pRN3-RFP vectors using the In-Fusion cloning kit (Clontech, Takara). *X. laevis cyclin B3* cDNA (clone ID: 781186, NCBI ID: 379048) was purchase from Dharmacon and used as a template for PCR amplification. *D. rerio* and *D. melanogaster cyclin B3* (NCBI ID: 767751 and 42971, respectively) were amplified by PCR from cDNA from *D. rerio* embryo or the S2 cell line, respectively. Amplified products for *X. laevis*, *D. rerio* and *D. melanogaster* were cloned into pRN3-GFP vector using the In-Fusion cloning kit (Clontech, Takara). To generate *Ccnb3* MRL mutation, overlapping primers with specific mutation were used to amplify PCR product from the wild-type template pRN3-cyclin B3-RFP, the template was digested by DpnI treatment, and the PCR product was used to transform *E. coli* DH5α competent cells. To generate the ΔD-box mutation, PCR primers that delete the D-box were used to amplify the *Ccnb3* plasmid, and the resulting PCR product was phosphorylated and ligated. Primer list can be found as **Table S1**. Plasmids for in vitro transcription of cyclin B1-GFP, securin-YFP, histone H2B-RFP, cyclin A2-GFP and β-tubulin-GFP have been described [31, 32, 69, 72, 73]

### Expression and purification of recombinant proteins

Vectors for expression of cyclin B3 (wild type or MRL mutant) N-terminally tagged with maltose-binding protein and 6x histidine were generated by cloning in pFastBac1-MBP6XHis. Wild-type *Ccnb3* was amplified from pRN3-cyclin B3-RFP and the MRL mutant was amplified from pRN3-*cyclin B3 MRL*-RFP. The plasmid was digested with SspI restriction endonucleases. By using In-Fusion cloning kit (Clontech, Takara) the PCR products were cloned into SspI linerazed plasmid. Viruses were produced by a Bac-to-Bac Baculovirus Expression System (Invitrogen) following the manufacturer’s instructions. Baculovirus expressing CDK1 and CDK1-HA were generously supplied by Dr. R. Fisher [75].

To express ^MBPHis^cyclin B3 and ^MBPHis^cyclin B3 MRL alone, *Spodoptera frugiperda* Sf9 cells were infected with virus at a multiplicity of infection (MOI) of 3. To express ^MBPHis^cyclin B3-CDK1or CDK1-HA and ^MBPHis^cyclin B3 MRL-CDK1or CDK1-HA complexes, Sf9 cells were infected with both viruses at a multiplicity of infection (MOI) of 3 and 2, respectively. Cells from two 150 mm plastic dishes per infection (each containing ~ 3×10^7^ cells) were harvested 48 hours after infection by centrifugation at 1500 rpm for 5 minutes and then washed with ice-cold PBS. Cell pellets were resuspended in 1.7 ml of ice-cold lysis buffer (25 mM HEPES-NaOH pH 7.5, 150 mM NaCl, 100 μM leupeptin (Sigma-Aldrich, L2884)), 1× Complete protease inhibitor tablet (Roche) and 0.5 mM phenylmethanesulfonyl fluoride (PMSF)). Cells were lysed by sonication and centrifuged at 15000 rpm for 30 minutes, at 4°C. The cleared extract was moved to a 2 ml Eppendorf tube and mixed with 300 µl slurry of amylose resin (NEB, E802L) pre-equilibrated with lysis buffer. After 1 hour at 4°C with end-over-end rotation, the amylose resin was centrifuged at 300 *g*, for 1 minute, at 4°C. Resin was washed briefly with ice-cold lysis buffer before transfer on Bio-Spin chromatography columns (BioRad, 7326008). The resin on the column was washed five times with ice-cold wash buffer (lysis buffer plus 10% glycerol). To elute the proteins from the column, the resin was incubated with 100–200 μl ice-cold elution buffer (wash buffer plus 10 mM maltose (Sigma, M5885)). All steps during the purification were done at 4°C.

### Histone H1 kinase assay with recombinant proteins expressed in Sf9 cells

Eluates from the amylose affinity purification step were used for histone H1 kinase assays. Reactions were performed in 10 mM HEPES-NaOH pH 7.4, 75 mM NaCl, 1 mM DTT, 10 mM MgCl_2_, 100 μM ATP (Roche, 11140965001), 1 μCi [γ–^32^P] ATP and histone H1 (5 μg/μl) (Sigma, 10223549001). 10 μl reaction mix was added to 10 μl eluate from the amylose resin and incubated for 30 minutes at room temperature. To terminate reactions, 5 μl of 4× SDS sample buffer was added to each reaction and samples were boiled for 5 mins before loading on 12% NuPage Bis-Tris NuPAGE precast gels (Life Technologies). Samples were subjected to electrophoresis at 150 V for 60 minutes. The gel was then vacuum dried for 45 minutes, at 80°C. Radiolabelled species were imaged using a Fuji phosphoimager and analyzed by Image Gauge software.

### Histone H1 kinase assays using oocyte extracts

Kinase assays to determine endogenous MPF activity were performed on aliquots of 5 oocytes at the indicated stages of meiotic maturation such as described before [30]. In short, oocytes were lysed in 3 μl of lysis buffer ((50 mM Tris (pH 7.5),150 mM NaCl,1% Igepal (Sigma),10% glycerol,2 mM EDTA, supplemented with protease inhibitors (Roche, Mini tablets) on ice for 20 minutes before adding 6 μl kinase assay buffer (50mM Tris (pH 7.5), 150 mM NaCl, 10mM MgCl_2_, 0.5 mM DTT, 2.5 mM EGTA, 150 μM ATP) supplemented with 3 μCi of [γ-^32^P]ATP (Perkin Elmer) per sample. Histone H1 (Millipore, 0.5 μg per sample) was used as a substrate. Kinase reactions were incubated 30 min at 30°C, denatured, and analyzed by SDS-PAGE. The gel was fixed and dried before being exposed and scanned to detect incorportation of [γ-^32^P]ATP into the substrate, using a Typhoon FLA9000 phosphorimager (GE Healthcare Life Sciences). Scans were analyzed by ImageJ.

### Microinjection and live imaging

*In vitro* transcription of all mRNAs was done using the Ambion mMessage Machine kit according to the manufacturer’s instructions. mRNAs (1 to 10 pM) were purified on RNeasy purification columns (Qiagen). GV stage oocytes were microinjected with mRNA on an inverted Nikon Eclipse Ti microscope. Microinjection pipettes were made using a magnetic puller (Narishige PN-30). Oocytes were manipulated with a holding pipette (VacuTip from Eppendorf) and injection was done using a FemtoJet Microinjector pump (Eppendorf) with continuous flow. Injection of the oocytes was done on a Tokai Hit temperature controlled glass plate at 38°C. Live imaging of oocytes in Figures 3B, 3E and 7B was carried out on an inverted Zeiss Axiovert 200M microscope coupled to an EMCCD camera (Evolve 512, Photometrics), combined with an MS-2000 automated stage (Applied Scientific Instrumentation), a Yokogawa CSU-X1 spinning disc and a nanopositioner MCL Nano-Drive and using a Plan-APO (40×/1.4 NA) oil objective (Zeiss). Alternatively, a Nikon eclipse TE 2000-E inverted microscope with motorized stage, equipped with PrecisExite High Power LED Fluorescence (LAM 1: 400/465, LAM 2: 585), a Märzhäuser Scanning Stage, a CoolSNAP HQ2 camera, and a Plan APO (20×/0.75 NA) objective was used. For Figures 3B, 3E and 7B, images of chromosomes were acquired using the inverted microscope Zeiss Axiovert 200M with 11 z-sections of 3 µm. Both microscopes were controlled by Metamorph software.

### Chromosome spreads and immunofluorescence

Oocytes were rinsed in successive drops of Tyrode’s acid solution to remove their *zona pellucida.* For chromosome spreads, oocytes were fixed at room temperature using a spread solution (1% paraformaldehyde, 0.15% Triton X-100 and 3 mM DTT, from Sigma-Aldrich), at the time points indicated. For whole-mount immunofluorescence, chambers were coated with concanavaline A (Sigma-Aldrich) in M2 PVP (polyvinylpyrrolidone 0,1mM from Merck-Millipore) medium and oocytes were placed in the chambers and centrifuged at 1400 rpm for 13 min at 38°C. Oocytes were then placed in a cold treatment solution (20 mM HEPES-NaOH, 1 mM MgCl2, pH 7.4) on top of ice (4°C) for 4 min to remove unstable microtubules. Following cold treatment, oocytes were incubated in a formaldehyde fixation solution (BRB80 medium with 0.3% Triton X-100, 1.9% formaldehyde (Sigma-Aldrich)) at 38°C for 30 min. Primary antibodies were used at the indicated concentrations: human CREST serum autoimmune antibody against centromere (Immunovision, HCT-0100, 1:100), rabbit polyclonal anti-MAD2 (1:50)[71], and mouse monoclonal anti-alpha-tubulin (DM1A) coupled to FITC (Sigma-Aldrich, F2168, 1:100). Secondary antibodies were used at the following concentrations: anti-human Alexa 488 (Life Technologies, A11013, 1:200), anti-human Cy3 (Jackson ImmunoResearch, #709-166-149, 1:200), anti-rabbit Cy3 (Jackson ImmunoResearch, #711-166-152, 1:200). Hoechst 3342 (Invitrogen, H21492) at 50 µg/ml was used to stain chromosomes and AF1 Cityfluor mounting medium was used.

### Image acquisitions of fixed oocytes

An inverted Zeiss Axiovert 200M microscope as described above was used to image chromosome spreads and for whole mount immunofluorescence acquisitions, using a 100×/1.4 NA oil objective coupled to an EMCCD camera (Evolve 512, Photometrics). Six z-sections of 0.4 µm were taken for spreads, and 11 z-sections of 1 µm for whole mount oocytes. Stacks were assembled in ImageJ.

### Quantification of fluorescent signals

For the quantification of MAD2 fluorescence signal, the intensity was calculated for each kinetochore using an 8 × 8 pixels square on the kinetochore (where CREST signal is located). The same sized square was used adjacent to the MAD2 fluorescence to measure background, which was subtracted from the MAD2 signal. The MAD2 fluorescence intensity was normalized to the CREST signal of the same kinetochore. For the cyclin B3-RFP, ΔDbox cyclin B3-RFP, cyclin A2-GFP, cyclin B1-GFP, and securin-YFP quantifications, a 150 × 150 pixels circle was placed in the center of each oocyte for the fluorescence intensity signal measurement, and another placed adjacent to the oocyte to measure background. For each oocyte, background-subtracted values were normalized relative to the highest value. All measurements were done using ImageJ software.

### Statistical analysis

For statistical analysis, GraphPad Prism 7 was used. For the comparison of independent samples, unpaired Student’s t test was carried out. Error bars indicate means ± standard deviation. Sample size and statistical tests are indicated in the figure and figure legends, respectively. Selection of oocytes for different conditions was at random. No statistical analysis was used to determine sample size. Collection and analysis of the data were not performed blind to the conditions of the experiments, no data from experiments were excluded from analysis.

## Supplemental Material

### Supplemental Movie legends

**Movie 1.** Progression through meiosis I in control oocyte. Representative movie of a *Ccnb3^+/–^* oocyte incubated with SiRDNA to visualize chromosomes. Acquisitions were started at GVBD + 6:30 (hours:minutes). Eleven z-sections with 3 μm steps were taken to follow chromosome movements, and shown are overlays of the stack. Acquisitions of the far-red and DIC channel were done every 20 min, 21 time points are shown. Related to Figure 2A. (n = 42)

**Movie 2.** Progression through meiosis I in *Ccnb3^−/−^* oocyte. Representative movie of a *Ccnb3^−/−^* oocyte incubated with SiRDNA to visualize chromosomes. Acquisitions were started at GVBD + 6:30 (hours:minutes). Eleven z-sections with 3 μm steps were taken to follow chromosome movements, and shown are overlays of the stack. Acquisitions of the far-red and DIC channel were done every 20 min, 21 time points are shown. Related to Figure 2A. (n = 63)

### Supplemental table legend

**Table S1.**Primers used for cloning expression plasmids. Primers used to generate the indicated plasmids are shown. See Methods section for details.

## End Matter

### Author Contributions and Notes

M.E.K. generated the *Ccnb3^−/−^* mouse strain and cloned plasmids used in this study, unless otherwise noted. Experiments in Figures 1A-C, 3C, and 6A were performed by M.E.K. Experiments in Figures 6B, C, and 7A, B were performed by N.B, and the kinase assays in Figure 3D by K.W. The remaining experiments and statistical analysis were done by N.B. and M.E.K. Overall supervision, funding acquisition and project administration was done by K.W and S.K.. Figures were prepared by N.B., M.E.K. and K.W., and the manuscript was written by K.W. and S.K. with substantial imput from all authors. The authors declare no conflict of interest.

## Acknowledgments

We thank W. Li, Q. Zhou, and Y. Zhang (State Key Laboratory of Stem Cell and Reproductive Biology, Institute of Zoology, Chinese Academy of Sciences) for sharing unpublished data. We thank A. Koff (MSKCC) for sharing unpublished data and for discussions and critical reading of the manuscript. We thank P. Romanienko, W. Mark, J. Ingenito and J. Giacalone (MSKCC Mouse Genetics Core Facility) for generating CRISPR/Cas9-targeted mice and J. White (MSKCC Laboratory of Comparative Pathology Core Facility) for anatomic pathology. We thank R. Fisher (Icahn School of Medicine at Mount Sinai) for sharing CDK1 and CDK1-HA carrying baculoviruses and advice for the CDK expression and histone H1 kinase assays. We thank C. Claeys Bouuaert (Keeney lab) for help with the baculovirus expression system and purification of cyclin B3-CDK1. We thank M. Boekhout and D. Ontoso (Keeney lab) for help with animal handling. We thank B. Joseph (MSKCC) for *Drosophila* cDNA, M. Jelcic and C. Huang (MSKCC) for *D. rerio* cDNA, and H. Funabiki (Rockefeller University) for *Xenopus* cDNA. We thank N. Fang, M. Turkekul, A. Barlas and K. Manova-Todorova (MSKCC Molecular Cytology Core Facility) for help with ovarian cytology. We thank members of the Wassmann and Keeney labs for discussion, S. Touati (Crick, London) and A. Karaiskou (CDR Saint Antoine, Paris) for comments on the manuscript, E. Nikalayevich (IBPS) for help with the separase sensor assay, F. Passarelli (IBPS) for help with rescue experiments in Figure 7, C. Rachez (Institut Pasteur, Paris) for access to the phosphorimager, and D. Cladière (IBPS) for technical help and help with animal handling. MSKCC core facilities were supported by NIH grant P30 CA008748. N. Bouftas received a 3-year PhD fellowship from the French Ministère de la Recherche. Work in the Wassmann lab was financed through a grant "Equipe FRM" by the Fondation de la Recherche Médicale (Equipe DEQ20160334921 to KW), an ANR grant (ANR-16-CE92-0007-01 to KW), as well as core funding from UPMC and the CNRS. Work in the Keeney lab was supported by the Howard Hughes Medical Institute.

